# Computing structure-based lipid accessibility of membrane proteins with mp_lipid_acc in RosettaMP

**DOI:** 10.1101/086579

**Authors:** Julia Koehler Leman, Sergey Lyskov, Richard Bonneau

**Affiliations:** Center for Computational Biology, Flatiron Institute, Simons Foundation, 160 Fifth Avenue, New York, NY 10010, USA; Departments of Biology and Computer Science, Center for Genomics and Systems Biology, New York University, New York, NY 10003, USA; Department of Chemical and Biomolecular Engineering, Johns Hopkins University, Baltimore, MD 21218, USA

**Keywords:** membrane proteins, structure prediction, lipid accessibility, accessible surface area, Rosetta software

## Abstract

**Background:** Membrane proteins are vastly underrepresented in structural databases, which has led to a lack of computational tools and the corresponding inappropriate use of tools designed for soluble proteins. For membrane proteins, lipid accessibility is an essential property. Even though programs are available for sequence-based prediction of lipid accessibility and structure-based identification of solvent-accessible surface area, the latter does not distinguish between water accessible and lipid accessible residues in membrane proteins.

**Results:** Here we present `mp_lipid_acc`, the first method to identify lipid accessible residues from the protein structure, implemented in the RosettaMP framework and available as a webserver. Our method uses protein structures transformed in membrane coordinates, for instance from PDBTM or OPM databases, and a defined membrane thickness to classify lipid accessibility of residues. `mp_lipid_acc` is applicable to both α-helical and β-barrel membrane proteins of diverse architectures with or without water-filled pores and uses a concave hull algorithm for classification. We further provide a manually curated benchmark dataset, on which our method achieves prediction accuracies of 90%.

**Conclusion:** We present a novel tool to classify lipid accessibility from the protein structure, which is applicable to proteins of diverse architectures and achieves prediction accuracies of 90% on a manually curated database. `mp_lipid_acc` is part of the Rosetta software suite, available at www.rosettacommons.org. The webserver is available at http://rosie.graylab.jhu.edu/mp_lipid_acc/submit and the benchmark dataset is available at http://tinyurl.com/mp-lipid-acc-dataset.

**Supplementary information:** Supplementary information is available at *BMC Bioinformatics*.

## 1 Background

Membrane proteins carry out a variety of essential functions and are targeted by over half of drugs in use [1], yet only make up ~2% of proteins in the Protein Data Bank due to difficulties in structure elucidation. The dearth of structures has in turn led to a lack of prediction tools, which have typically focused on either α-helical or β-barrel membrane proteins [2], or on specific features like the prediction of membrane pores [3]. One important characteristic of membrane proteins is accessibility to the lipid environment of the bilayer. Knowledge of lipid accessibility is important for our understanding of membrane protein structure, stability [4], interactions inside the bilayer, the development of membrane protein energy functions, the development of sequence-based predictors, and as an indicator of binding interfaces. While sequence motifs for helix-helix interactions in the membrane have been well-studied [5], the broader picture of large protein-protein interfaces in the membrane is still developing.

Prediction of lipid accessibility is not trivial: the protein ‘interior’ can either be water accessible in case of pores, or buried hydrophobic residues. For the latter, the hydrophobicity profile of buried residues is similar to lipid-facing residues, hence hydrophobicity as a solitary feature would be insufficient for classification. Solvent accessible surface area (SASA) is useful to identify accessibility of the residue to the solvent, yet it does not distinguish between lipid or water accessibility. Further, geometric considerations might be useful for the structures of β-barrels, as sidechains typically either face into the barrel or away from it, but this distinction is typically less clear for α-helical proteins. Therefore, a combination of different features is required for accurate prediction of lipid accessible residues.

The problem is further complicated by the fact that accessibility to lipid relies on protein embedding in the membrane and experimental measurements of membrane embedding features (like depth, angle, and membrane thickness) are challenging. Lipid molecules are only present in a few crystal structures, and the model membrane used for crystallization or structure determination is very different from a native membrane environment because it lacks the exact lipid composition [6], membrane thickness, asymmetry [7], lateral pressure [8] and shape [9] (compare micelles, bicelles, nanodiscs, lipidic cubic phases vs. a flat or curved bilayer). Protein embedding in the membrane is therefore predicted using computational approaches that rely on different score functions, such as TMDET [10], PPM [11], and iMembrane [12]. However, benchmarking of these tools is difficult without high-resolution experimental data. Similarly, for lipid accessibility there are no compiled structural databases to test the performance of a predictor from the protein structure, hence expert manual curation of such a database is required.

A range of **sequence-based predictors** is available that specializes either on transmembrane span or topology prediction for α-helical bundles or β-barrels, SASA, or lipid accessibility. Trans-membrane span predictors classify residues as *in the membrane* vs. *in solution* and are able to achieve prediction accuracies >90% in the two-state scenario [13], [14]. Topology predictors [15], [16] classify residues as *inside*/*outside* the cell or organelle and *in the membrane* (for definitions for the OPM (Orientations of Proteins in Membranes) database [11], see http://opm.phar.umich.edu/about.php?subject=topology). SASA predictors classify residues as *buried* or *exposed* to the solvent without specifying whether the solvent is water or lipid. They typically use support vector machine approaches and report prediction accuracies in the range of 70-75%: the ASAP tool can be used for both α-helical and β-barrel membrane proteins [17], the MPRAP method is optimized for SASA prediction of both soluble and transmembrane regions of membrane proteins [18], and the TMExpo tool predicts embedding angles in addition to SASA [19]. Further, pore-lining residues and channels can be predicted from the protein sequence using the recently developed tool PRIMSIPLR [3], which achieves a prediction accuracy around 86%. Lipid accessibility (*lipid exposed* versus *lipid buried*) can be predicted from the protein sequence with the LIPS server [20], which uses contact maps between helical faces and residue conservation for classification. It achieves a prediction accuracy of 88%, according to the authors.

**SASA prediction from the protein structure** can be obtained with a few early predictors (Naccess [21], MSMS [22], GetArea: http://curie.utmb.edu/getarea.html [23]); unfortunately, some of these methods lack benchmarking results, were only tested on a few low-resolution crystal structures or lack availability as a web server and/or documentation. The recently developed, well-documented 3V web interface for volume, solvent exclusion and channel prediction does not compute SASA values per residue [24].

Here, we present a method to identify lipid accessible residues from a protein structure that is already embedded in membrane coordinates. Our method uses a 2D concave hull algorithm on a point cloud generated by projecting Cα coordinates from horizontal slices of the protein within the bilayer onto the plane of the membrane. For each slice, the convex hull, the concave hull, and a ‘concave shell’ is computed, through which lipid accessible residues are identified (see Implementation and Fig. 2). The classification of the concave shell is output as adjusted B-factors in the PDB file, which can be easily visualized or extracted for further analysis. We compute prediction accuracies on a manually curated benchmark dataset available for download at http://tinyurl.com/mp-lipid-acc-dataset. Our method `mp_lipid_acc` is also publicly available as a webserver on ROSIE [25] at http://rosie.graylab.jhu.edu/mp_lipid_acc/submit. To our knowledge, this is the first tool to classify lipid exposure of residues from a protein structure. Our method is applicable to both α-helical and β-barrel membrane proteins of any architecture and will be useful for a variety of problems from score function derivation, development of sequence-based predictors, and structural modeling.

**Figure 1:**
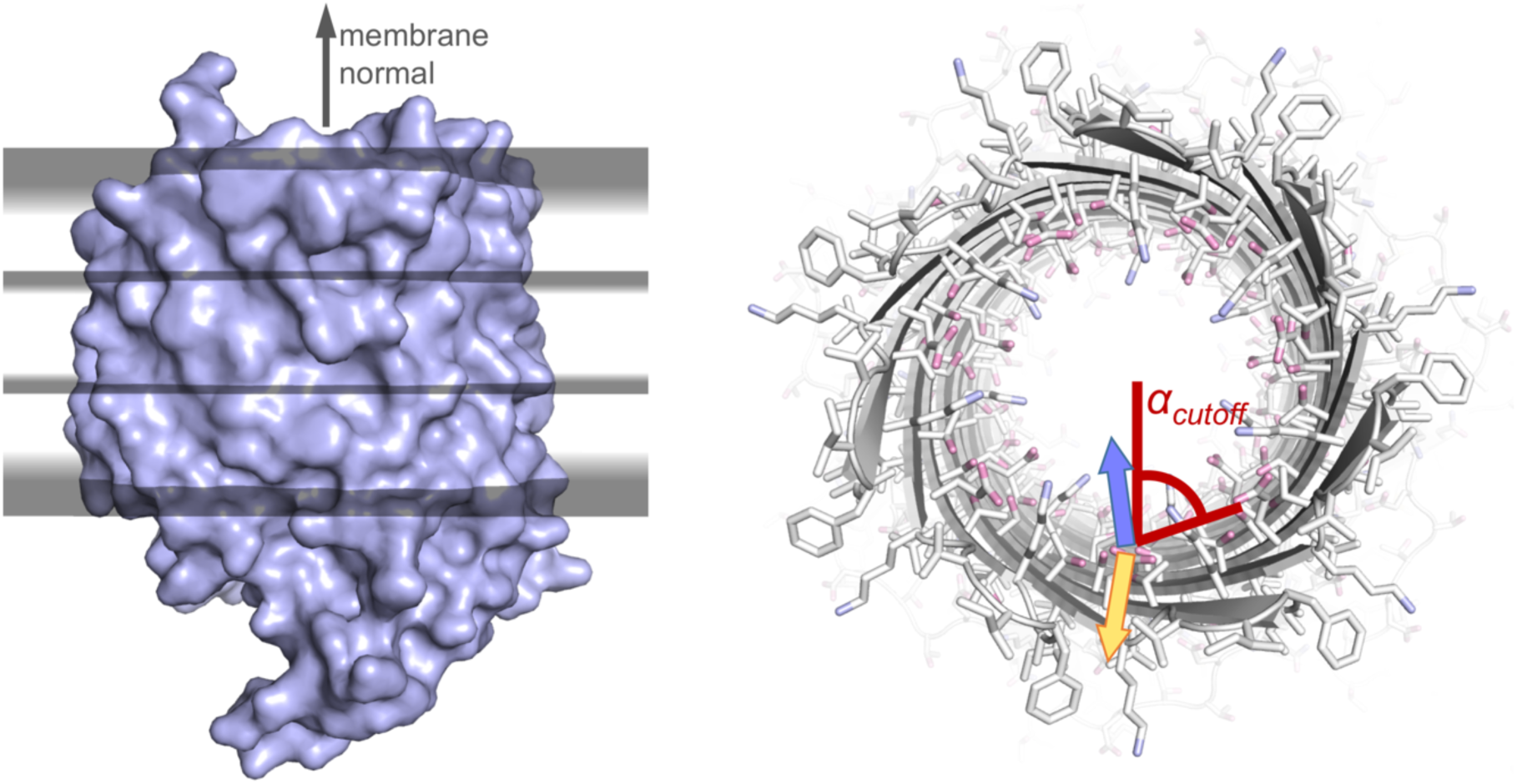
Algorithm. In our algorithm `mp_lipid_acc`, the protein is cut into horizontal slices (left) and the center-of-mass in each slice is computed from the coordinates of the Cα atoms. In each slice, the coordinates of the Cα atoms are projected onto the xy-plane, from which points first the convex hull, then the concave hull, and then the ‘concave shell’ are computed (see Implementation). For β-barrels, only residues which are part of the concave shell and have a Cα-Cβ-COM angle larger than a cutoff value of 65 degrees are classified as lipid accessible (right). For instance, the residue with a sidechain orientation represented by the yellow vector in the panel on the right would be lipid accessible, whereas the sidechain in blue would be lipid inaccessible.

**Figure 2:**
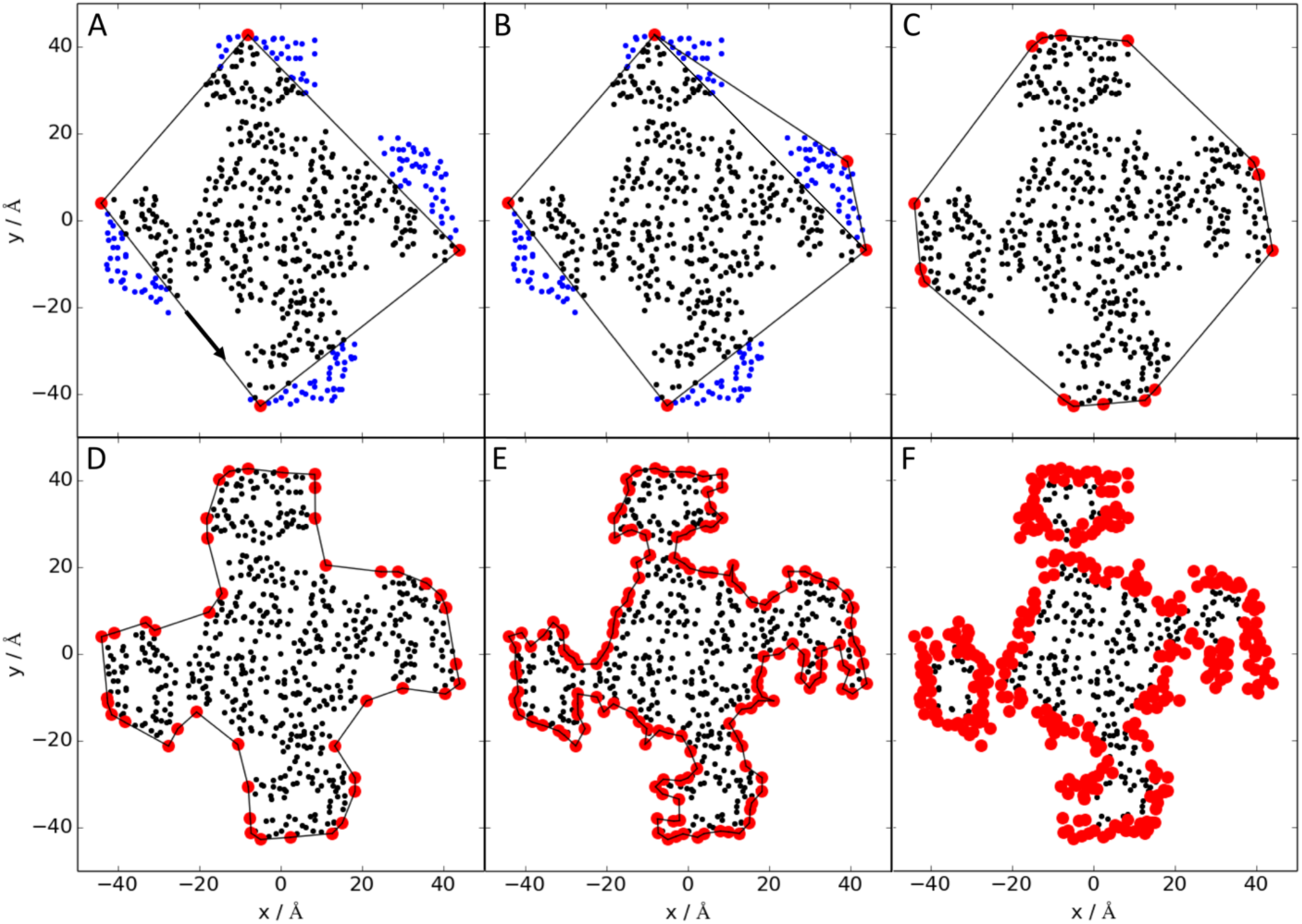
Convex and concave hull algorithms. Illustration of the convex hull, concave hull, and ‘concave shell’ from a 2D point cloud that was projected onto the xy plane from the 3D structure. The example protein is the sodium channel with PDBID 4DXW and the points in blue are outside the hull, the points in black are inside the hull and the points in red are part of the hull. The convex hull algorithm starts by connecting the points with smallest and largest x and y values in counter-clockwise direction and identifying the points inside the hull (A). The hull is extended to a convex hull by connecting two points on the boundary, finding the point in the outside list that is farthest away from the two boundary points and that is in clockwise direction, and adding this point to the boundary (B). The points within the triangle between the old two boundary points and the added one are then classified as inside the hull. These steps are repeated until all points are inside the convex hull (C). The concave hull is then computed by finding the longest distance between two connected points on the boundary, finding the point inside the hull that has the smallest enclosing angles with respect to the two original boundary points, and adding this new point to the hull boundary. The distance cutoff defines the ‘resolution’ of the hull with a distance cutoff of 15 Å in (D) and 5 Å in (E). This process is repeated until all distances between connecting points are smaller than the cutoff distance. Lastly, a concave shell is computed by including points within an xy distance radius from the original boundary points (from E to F). All boundary points in F are now part of the concave shell and classified as lipid exposed.

## 2 Implementation

### 2.1 Curation of training and benchmark datasets

We created a small dataset of 14 proteins for the development of the algorithm (Supplementary Table 1) and to find an optimal parameter set. These proteins cover a wide range of protein folds from one or two transmembrane helices, helical bundles with and without smaller pores (for instance GPCRs, aquaporins), small and large helical pores, transporters (ABC transporter), ion channels with voltage sensor domains, oblong helical channels (for instance chloride channel), β-barrels with and without internal domains, β-barrel monomers and trimers, and cigar shaped β-barrels and pore-forming toxins. The PDBIDs for this dataset were 1afo, 1ek9, 1fep, 1kpk, 1qd6, 2kix, 2r9r, 2rh1, 2wcd, 3emn, 3ne2, 3wmf, 4tnw, 7ahl.

We curated a benchmark dataset for testing our method (Supplementary Table 2): all membrane protein chains were downloaded from the PDBTM database [26], which were then culled with the PISCES server [27] with the following parameters: sequence identity ≤ 25%, resolution ≤ 3.0 Å, R-factor cutoff 0.3, sequence length 40-10,000, include non-Xray entries, exclude Cα-only entries, cull by chain. We then excluded all EM structures with resolutions > 3 Å and removed all XFEL structures with resolution “0.0” or proteins for which no method was specified. Further, we removed photosynthetic reaction centers and photosystems I and II as they have very loosely packed helices complexed with a large number of interstitial chlorophyll molecules. We also removed the proteins present in the training dataset.

The structures in each set were downloaded from the PDBTM database [26], for which the proteins are already transformed into the membrane coordinate frame: the membrane center is defined as the origin at (x, y, z) = (0, 0, 0) and the membrane normal vector lies along the z-axis with (0, 0, 1). To compute the membrane embedding of a protein structure, PDBTM uses the TMDET algorithm [10], [28] that fits the membrane embedding according to an objective function that contains measures of hydrophobicity and structural information regarding the Cα-trace within a protein slice, such as straightness, turns and termini. Initially, we downloaded structures from the OPM database [11], but through visual inspection of the entire database we realized that the protein embedding seems generally better in PDBTM. The structures were then cleaned from additional atoms (such as ligands and co-factors), renumbered, and the span files were computed as described previously [29] with a fixed membrane thickness of 30 Å. We used a fixed membrane thickness to avoid a circular influence of the thickness prediction from PDBTM onto our benchmark dataset.

We ran our algorithm over the benchmark dataset (204 proteins) as a first pass, which outputs the lipid accessibility as a modified B-factor column in the PDB file. We then manually corrected classification errors by visualizing each protein in PyMOL [30], coloring it according to B-factors, and then manually adjusting the B-factors for each incorrectly classified residue. Our benchmark dataset is described in Supplementary Table 2 and is available for download at http://tinyurl.com/mp-lipid-acc-dataset.

### 2.2 Algorithm overview

The protein was cut into 10 Å thick horizontal slices along the membrane normal (Fig. 1). For each slice, the Cα atoms were projected onto the xy plane, from which points the convex hull was computed. The convex hull is the smallest set of points on the outer perimeter of the 2D point cloud that encloses the entire point cloud; it is computed by a *QuickHull* algorithm [31], [32] (see below). From the convex hull, we computed the concave hull [33], which defines the set of points on the outer perimeter of the point cloud. It has a smaller surface area than the convex hull and encloses the point cloud by concave surfaces (see below). We then computed the concave shell that includes points (Cα atoms) whose xy coordinates are within a radius of the original boundary points. Concave shells were computed for each horizontal slice in the membrane region.

Residues with Cα atoms in the concave shell are classified as lipid exposed with the following exceptions: (1) if the number of TM spans is smaller or equal 2, *all* residues in the membrane region are classified as lipid exposed; (2) for β-barrels, we make the distinction of whether the sidechain faces into the interior of the barrel or not. For each slice, we therefore compute the center-of-mass (COM) of the Cα atoms and only classify residues with the Cβ-Cα-COM angle larger than 65 degrees as lipid accessible (for Glycine, we used 2HA to represent ‘Cβ’). (3) For smaller helical bundles with 7 or less TM spans, the concave shell algorithm over-predicted lipid accessible residues in the interior of the protein. To counter balance this over-prediction, we took into account an angle cutoff of 45 degrees as described above.

### 2.3 Convex and concave hull

The **convex hull algorithm** is a classification of a 2D point cloud into three lists: inside the hull, outside the hull, and part of the hull (i.e. on the boundary). The *Quickhull* algorithm starts by classifying all points as outside the hull [31], [32]. Then, the points with the smallest x value, smallest y, largest x and largest y are connected and moved to the list of points on the boundary. The points inside the rectangle are moved to the inside list (Fig. 2A). Then, two neighboring points in the boundary list are connected by a line and the point outside with the largest distance in clockwise direction is moved to the boundary (Fig. 2B), while the points inside this triangle are added to the point list inside the hull. This last step is repeated until all points are either inside the hull or on the boundary and the outside list is empty. The convex hull algorithm identifies the points on the hull that make up a convex shape (Fig. 2C). However, it fails to identify points on the hull that encompass the smallest surface area.

The **concave hull** on the other hand, defines points with the smallest surface area, which requires ‘cutting into’ the boundaries of the convex hull to create concave surfaces. Starting from the convex hull, two neighboring points on the boundary are connected by a line [33]. If the distance of the line segment is larger than a pre-defined distance cutoff, the point inside the hull (counter-clockwise) is identified that has the smallest sum of angles to the two original boundary points. This point is added to the boundary and this last step is repeated until all the distances between neighboring points in the boundary list are smaller than the pre-defined cutoff. The cutoff is required because there is no unique solution to the classification problem as to which points are part of the concave hull. The cutoff is therefore a measure similar to a ‘resolution’ that defines how rugged the concave surfaces are (compare Fig. 2D and 2E).

To extend the information of the concave hull back into three dimensions, we classified points (i.e. Cα atoms) within a certain xy-distance from the points on the boundary as part of the boundary – we call this the **concave shell** (Fig. 2F).

### 2.4 Adjustable parameters in the algorithm

Optionally adjustable parameters for this application are: (1) the width of the horizontal slices for which the concave shells are computed. As only Cα atoms are considered and to avoid overfitting of the convex shell due to data sparsity, the default slice width to is set 10 Å, corresponding to approximately 1/3 of the thickness of a physical membrane bilayer. The three slices therefore extend over the inner and outer leaflets and the space in between. Further, the membrane thickness should be an integer multiple of the slice width to avoid data sparsity in the last slice. The current membrane thickness is fixed at 30 Å. (2) The ‘resolution’ of the concave hull is defined by the distance cutoff between points on the hull boundary. 2D line segments between neighboring points on the hull are ‘cut in’ if their distance is longer than the distance cutoff (Fig. 2D and E). The default value for the distance cutoff is 10 Å, which is approximately the distance between two Cα atoms on the same side of neighboring helices. (3) To map the 2D concave hull back into three dimensions, Cα atoms with xy coordinates within a certain radius (shell radius) of the original boundary points are added to the hull, which we call the concave shell. The default shell radius is 6 Å, which is about half the diameter of an α-helix. (4) To distinguish between sidechains facing water-filled interiors in β-barrels from sidechains facing the lipid environment, we defined a cutoff for the Cβ-Cα-COM angle. The default value for the angle cutoff is 65 degrees, which is empirically determined and slightly smaller than 90 degrees to account for the curvature of β-barrels. (5) To classify lipid accessibility we distinguish between an α-helical bundle or β-barrel membrane protein, which refers to the secondary structure facing the lipid (i.e. barrel interior secondary structure is not considered). The type is auto-detected based on the prevalent secondary structure in the membrane, as computed by DSSP [34]. In the rare scenario that auto-detection fails, the secondary structure type can be set by the user.

If a residue is classified as lipid accessible, its B-factor is set to 50, otherwise it is set to 0. The PDB structure with the adjusted B-factor is output and can be visualized in PyMOL (The PyMOL Molecular Graphics System, Version 1.8 Schrödinger, LLC) with the provided script `color_bfactor.pml`.

## 3 Results and Discussion

We present an algorithm that classifies lipid exposed residues in membrane protein structures. It is applicable to monomeric and oligomeric α-helical and β-barrel membrane proteins with and without pores. Our algorithm uses protein structures transformed into membrane coordinates and a fixed membrane thickness and is applicable to a wide range of protein architectures. Classification is achieved through a 2D concave hull algorithm applied to the point cloud of membrane embedded Cα atoms projected onto the membrane plane. `mp_lipid_acc` then creates a PDB structure file with modified B-factors that can then be visualized by a provided PyMOL script. We further provide a publicly accessible webserver to run the classification, which is implemented in the ROSIE environment [25], accessible at http://rosie.graylab.jhu.edu/mp_lipid_acc/submit. Additionally, we make our manually curated benchmark dataset available to the public (http://tinyurl.com/mp-lipid-acc-dataset), which will be useful for the development of sequence-based predictors and membrane protein scoring functions.

The prediction accuracies for our benchmark set are shown in Table 1 with counts for true positives, true negatives, false positives, and false negatives. Accuracy, sensitivity and specificity are also computed. As a baseline for comparison, we used the prediction of rASA with a cutoff of 0.2. Table 1 shows that while rASA achieves almost as high prediction accuracies as our predictor, it is unable to distinguish between lipid accessible and water accessible residues, as demonstrated by the low sensitivity. In contrast, our simple algorithm correctly identifies lipid exposed residues, achieving a prediction accuracy of 91.2% (Table 1, Fig. 3, Table 2 and Supplementary Table 2), with consistently high sensitivity and specificity (~90%).

**Table 1.** Prediction accuracies in percent for the benchmark set. The first row indicates a prediction using relative accessible surface area in the membrane (cutoff 0.2) without lipid accessibility classification. The second row shows results for predicted lipid accessibility with our concave hull algorithm. Note that `mp_lipid_acc` is able to identify lipid exposed residues, giving rise to an almost 20% increase in sensitivity over a standard rASA algorithm.

|  | TP | TN | FP | FN | acc | sens | spec |
| --- | --- | --- | --- | --- | --- | --- | --- |
| rASA | 26526 | 119406 | 7727 | 10602 | 88.8 | 71.4 | 93.9 |
| hull | 33191 | 116548 | 10585 | 3937 | 91.2 | 89.4 | 91.7 |

TP: number of residues predicted as true positives; TN: true negatives; FP: false positives; FN: false negatives; acc: accuracy = $(TP+TN)/(TP+TN+FP+FN)$ ; sens = $TP/(TP+FN)$ ; spec = $TN/(TN+FP)$ .

**Figure 3:**
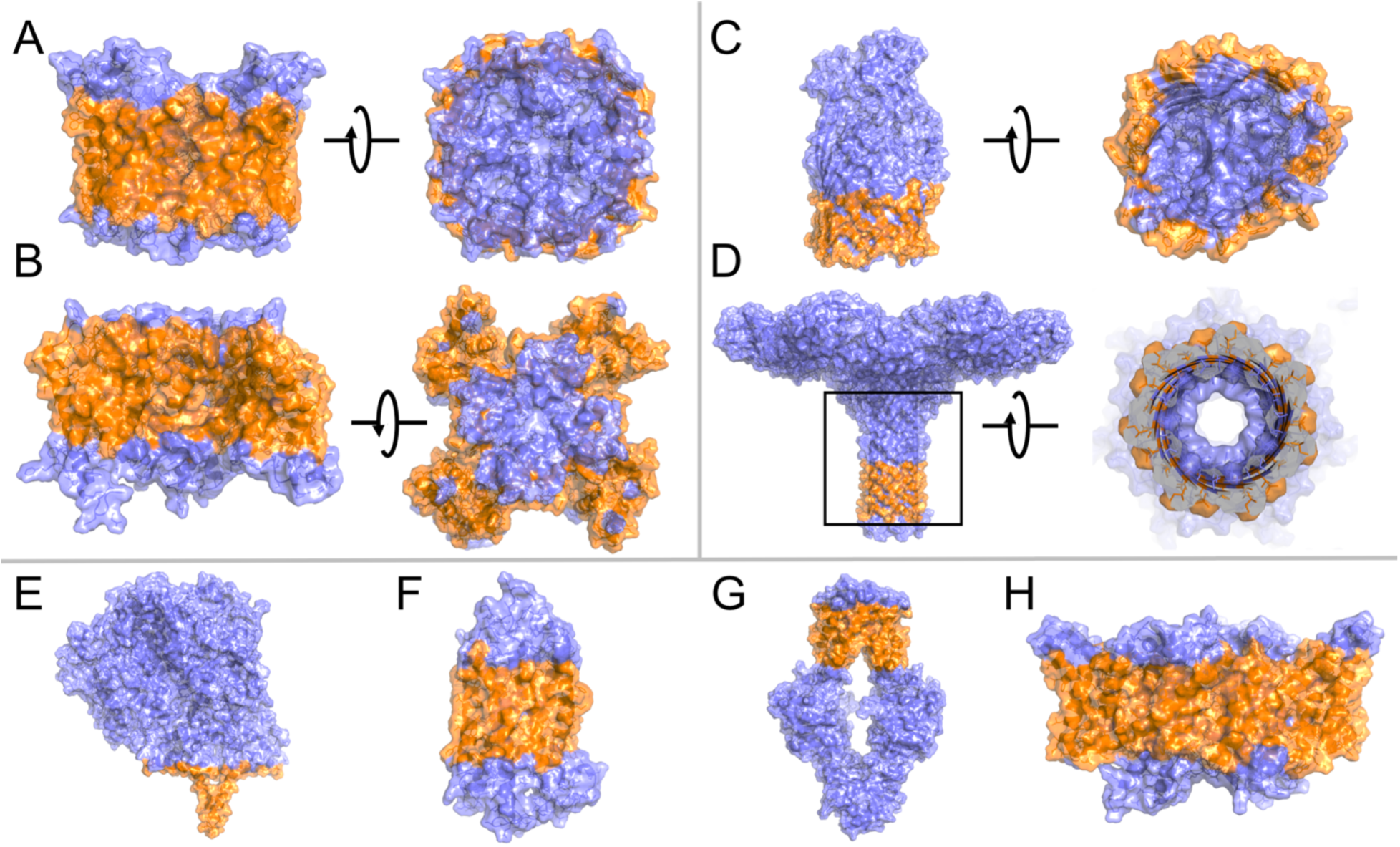
Classifications on selected examples from the benchmark dataset. Classifications for proteins of diverse architectures. Details about the individual proteins are provided in Table 2. In panel D, the pore of the protein (framed) is shown on the right, where the residues that protrude into solution, are clipped to show the prediction for the pore residues.

**Table 2.** Proteins for which predictions are shown in Fig. 3. A/B denotes α-helical vs. β-barrel membrane proteins. The benchmark dataset is shown in the Supplementary Material.

| PDBID | A/B | feature | family | subfigure |
| --- | --- | --- | --- | --- |
| 3D9S | A | aquaporin | aquaporin | A |
| 4DXW | A | ion channel | sodium channel | B |
| 3CSL | B | filled barrel | heme binding | C |
| 3W9T | B | pore-forming toxin | lectin | D |
| 2J7A | A | 1-2TM | nitrite reductase | E |
| 1U19 | A | GPCR | bovine rhodopsin | F |
| 3TUI | A | ABC transporter | methionine transporter | G |
| 4ENE | A | CIC channel | chloride channel | H |

We aimed to classify residues in well-packed α-helical proteins, helical pores, plain β-barrels, β-barrels with plug domains, monomeric and multimeric membrane proteins with and without pores, and multimeric ion channels with a large lipid exposed surface area due to their voltage sensor domains (Fig. 3). Because we are using a concave hull algorithm, `mp_lipid_acc` is applicable to a wide range of protein architectures that have perimeters of different shapes, for instance oblong protein structures (such as the chloride channel in Fig. 3H), ‘winged’ structures (such as ion channels with voltage sensor domains in Fig. 3B), and multimeric proteins.

Runtimes for `mp_lipid_acc` are consistently short, typically under 1 minute for proteins up to 2,000 residues, using default parameters (Fig. 4). The algorithm only takes into account Cα atoms in the protein. We tested using all atoms for the calculation of the hulls and the concave shell, which increased the runtimes considerably (over 1 hour), especially for large proteins, while achieving similar results. We therefore only make the option available to use Cα atoms.

**Figure 4:**
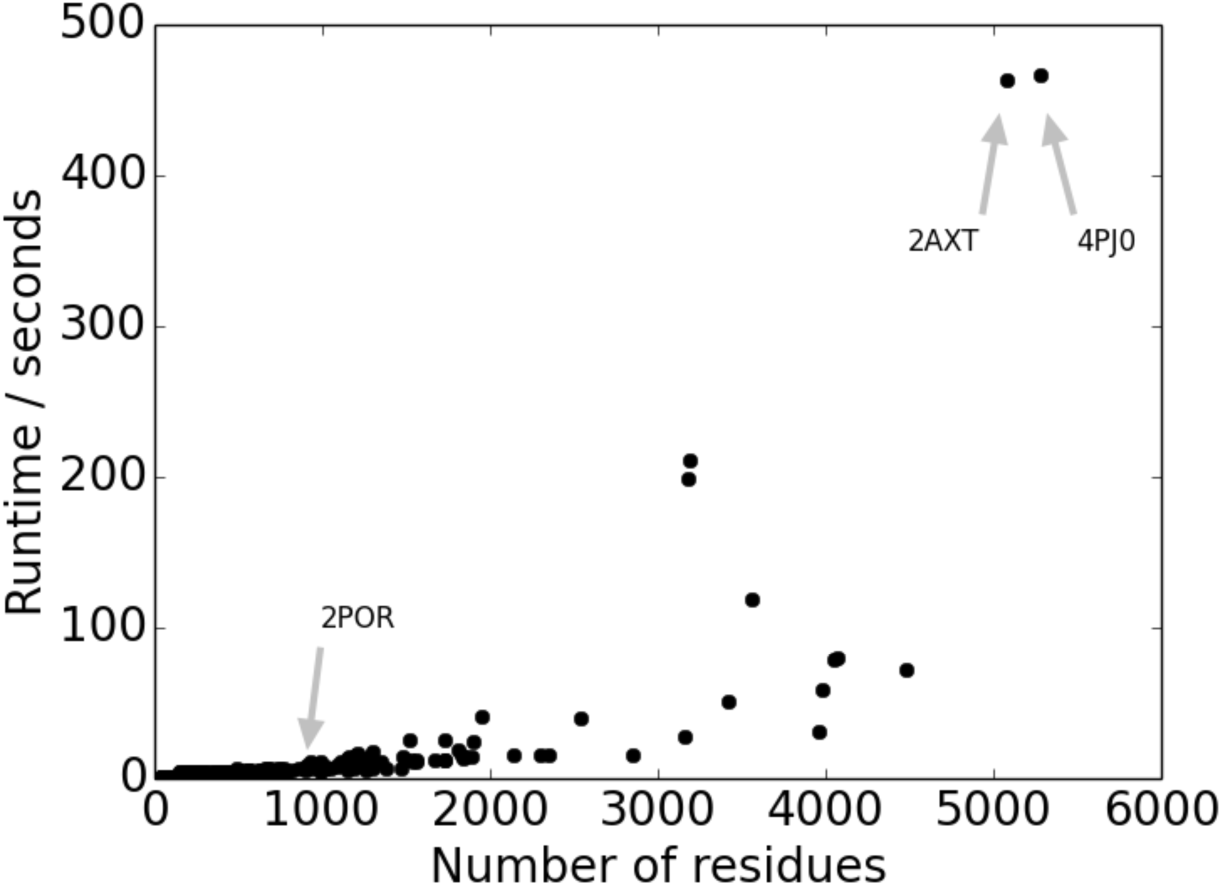
Runtimes for proteins of different sizes. Runtime of `mp_lipid_acc` in seconds for proteins of varying sizes. Runtimes are obtained when using the default parameters of a slice width of 10 Å, a distance cutoff of 10 Å, and a shell radius of 6 Å.

While improvements to the algorithm might lead to a smaller number of false positives, our algorithm is the first to classify lipid exposed residues from the protein structure, yielding prediction accuracies around 90%. It will be useful for molecular modeling and developing score functions and more sophisticated sequence-based approaches for the prediction of lipid accessibility. Classification of lipid accessibility from the structure is also useful to advance our understanding of membrane protein folding, stability and their interactions in the membrane bilayer [4].

## 4 Conclusion

Here we present a novel method to classify lipid accessibility in membrane protein structures. Our algorithm is applicable to α-helical and β-barrel membrane proteins with and without pores and for diverse protein architectures. `mp_lipid_acc` is implemented in RosettaMP and uses membrane embedded structures (from PDBTM or OPM) and a fixed membrane thickness to classify residues based on a concave hull algorithm. To test our method, we manually curated a benchmark dataset, on which `mp_lipid_acc` achieves accuracies, specificities and sensitivities around 90%. We believe that `mp_lipid_acc` and our benchmark set are an important first step for sequence- and structure-based prediction of lipid accessibility, score function optimization, and membrane protein modeling in general.

## Acknowledgements

The authors gratefully acknowledge Dr. Evan Baugh (NYU Biology dept.) for providing PyMOL/Python scripts used for the manual curation of the benchmark dataset.

